# Phenotypic and genotypic detection of the virulence factors and their association with antibiotic resistance in *Enterococcus* species

**DOI:** 10.1101/2023.04.03.535503

**Authors:** Tahani Momotaz, Fatima Afroz, Sharmin Chowdhury, Nahidul islam, Mohammad Tanvir Sarwar, Rehana Razzak Khan, Abu Naser Ibne Sattar

## Abstract

**Abstract-** Along with the emergence of drug resistant *Enterococcal* infection, role of various virulence factors in *Enterococci* is an emerging concept. A number of virulence factors like biofilm formation, hemolysin production, gelatin hydrolysis have important role in the pathogenesis of *Enterococci* and also associated with antibiotic resistance. The aim of our study was to detect the virulence factors and their encoding genes (*asa, gelE, esp, ebpR, hyl* gene for biofilm; *cylA* gene for hemolysis; *gelE* gene for gelatin hydrolysis) and also observe their association with antimicrobial resistance *Enterococci*. A total of 87 *Enterococci* were collected from different clinical samples. Virulence factors were detected phenotypically and antibiotic sensivity were done by Kirby Bauer disc diffusion method. Virulence genes were detected by conventional multiplex PCR and only the *ebpR* gene was detected by single conventional PCR. Majority of the isolated *Enterococci* were *E. faecalis* (75%) followed by *E. faecium* (23%) and (2%) *E. raffinosus* were also detected. About 52.3% of *E. faecalis* and 35% of *E. faecium* isolates were biofilm producers. Significant association was found between biofilm formation and *asa, esp, ebpR* genes both in *E. faecalis* and in *E. faecium.* Hemolysis was observed phenotypically in 30.8% isolates of *E. faecalis* and 20% isolates of *E. faecium*. Significant association was observed between *cylA* gene and hemolysin production in *E. faecalis*. Antibiotic resistance were higher in biofilm and hemolysin producing isolates of both species. Resistance to some antibiotics including ampicillin, ciprofloxacin, gentamicin were significantly higher among biofilm and hemolysin producer in *E. faecalis*.

## INTRODUCTION

*Enterococci* are member of normal gut flora of humans and animals that causes various community and hospital-acquired infections such as urinary tract infections, bacteremia endocarditis, meningitis and intra-abdominal infections [1].

The capacity of *Enterococc*i to acquire and share extra chromosomal elements including antibiotic resistance genes or virulence traits explains their increasing importance as nosocomial pathogens [2-4]. It is believed that virulence factors of *Enterococci* might have a role in increasing their capacity to colonize hospitalized patients [3]. The ability of biofilms formation is one of the important virulent characteristics of *Enterococci* [5]. Biofilms protect the bacteria from host immunological responses, phagocytosis and antibiotics [6], Numerous Enterococcal virulence factors including adhesions and secrated factors have been detected as being associated with the biofilm formation. Asa 1 (aggregation substance), Esp (extracellular surface protein) and Ebp (endocarditis and biofilm-associated pili) are the most important adhesion factors [7]. Asa1 is a pheromone-inducible protein that increases bacterial adherence to renal tubular cells [8]. Esp is a cell wall-associated protein that is responsible for colonization, persistence and biofilms formation in the urinary system [9]. The Ebp operon are associated with pilli and biofilm formation which is essential for causing UTI [8, 10]. For expression of Ebp operon, the *ebpR* (endocarditis and biofilm associated pilli regulator) gene is required and biofilm formation is reduced in *ebp* mutant strain [11]. Several secrated virulence factors of *Enterococci* including CylA (cytolysin), GelEA (gelatinase), and Hyl (hyaluronidase) play a role in pathogenesis of *Enterococci* [12]. Gelatinase is an extracellular zinc-containing metalloproteinase, the main role of gelatinase is providing nutrients to the bacteria by degrading host tissue. It has also some role in biofilm formation [13, 14]. Hyl acts on hyaluronic acid and consist of degradative enzyme that is responsible for tissue damage and facilitates spread of *Enterococci* through host tissue [15]. Cytolysin is a secreted toxin produced in response to pheromones, helps in the pathogenesis of *E. faecalis* by causing blood hemolysis [16].

Knowledge of the virulence factors of *Enterococci* may help to understand the pathogenic process of this microorganisms. So, this study was designed to detect different virulence factors and their encoding genes and their association with antibiotics resistance.

## MATERIALS AND METHODS

### Bacteria isolates

This cross sectional study was conducted at Department of Microbiology and Immunology, BSMMU over a period of 1 year from March 2019 to February 2020. A total of 87 *Enterococci* isolates were studied collected from different clinical samples (urine, blood, wound swab, pus and bile). *Enterococci* were identified in the laboratory by Gram stain, culture and standard biochemical test [17, 18].

### Antibiotic susceptibility test by disc diffusion

The Kirby-Bauer disk diffusion method was used for antimicrobial susceptibility testing of the isolated *Enterococci* using Mueller-Hinton agar and commercially available antibiotic disc (Biomaxima, Poland). Antibiotic discs of ampicillin (10 μg), cotrimoxazole (1.25/23.75 μg), ciprofloxacin (5 μg), gentamicin (120 (120 μg), nitrofurantoin (300 μg), vancomycin (30μg), linezolid (30 μg), teicoplanin (30 μg), fosfomycin (200 μg), quinopristin-dalfopristin (15 μg) were used. The disc concentration and zone of inhibition was used as recommended by the Clinical Laboratory Standards Institute guideline (CLSI, 2019) [19]. *S. aureus* ATCC 25923 was used as a standard strain.

### Phenotypic detection of the virulence factors

#### Detection of hemolysis

The hemolytic activity was detected by inoculating *Enterococci* on blood agar media. Hemolytic zone appears around the colony after 24 hours incubation at 37°C [20].

#### Gelatin hydrolysis activity

Gelatin production was detected by inoculating *Enterococci* on freshly prepared peptone yeast extract agar containing 4% gelatin and was incubated at 37°C for 24 hours and was cooled to room temperature for 2 hours. Appearance of turbid halo around the colonies indicates gelatin hydrolysis [21].

#### Study of Biofilm formation

Biofilm formation of *Enterococci* was done by tissue culture plate method (TCP) according to Toledo-Arana *et al* [22]. *Enterococci* were grown in Brain heart infusion broth (BHIB) (Becton Dickinson and company, USA) with0.25% glucose at 37°C overnight. The broth culture was diluted at a ratio of 1:40. 200μl of diluted culture suspension was inoculated in a sterile 96 well flat bottom polystyrene microtiter plate (Greiner Bio-One International, Kremsmunster, Austria). The positive control (*Klebsiella pneunoniae* ATCC 700603) and negative control (sterile BHIB-0.25% glucose) were also added in the same way. After overnight incubation at 37°C, the wells were washed with 200 μl of phosphate buffer saline (PBS) three times. The plate was air dried, fixed with 200 μl/well of 2% formalin at 4°C for 1 hour. Then stained with 1% crystal violet for 15 min and were rinsed under running tap water to remove the excess stain. Afterthat 200 μl ethanol acetone (80:20, v/v) was added in each well to solubilize crystal violet. Each assay was performed in triplicate and repeated three times. The optical density (OD) at 630 nm was measured using ELISA plate reader (Plate reader, model–A4, serial no.-1910, Das, Italy). The cut-off value (ODc) was calculated for each microtiter plate. ODc was of three standard deviations (SD) above the mean OD of the negative control: ODc = average OD of negative control + (3×SD of negative control). Final OD value of a tested strain was expressed as average OD value of the strain reduced by ODc value (OD = average OD of a strain - ODc). If a negative value is obtained, it should be presented as zero, while any positive value indicates biofilm production [23].

### Genotypic detection of virulence genes by PCR

DNA was extracted by boiling method [24]. Conventional multiplex PCR was performed to detect all virulence genes and single PCR were used to detect the *ebpR* gene. The PCR assay was performed in a reaction mixture with total volume of 25 μl containing 15 μl of master mix (HELINI, India), 0.15 μl Taq polymerase (Solis BioDyne Germany), 1 μl of forward and reverse primer each (10 pmol/μl), 3 μl of distilled water and 5 μl of undiluted extracted DNA. For all virulence genes the amplification condition was: pre-degeneration at 95 °C for 15 min followed by 30 cycles of degeneration at 94 °C for 1 min, annealing at 56 °C for 1 min and extension at 72 °C for 1 min and final extension at 72 °C for 10 min. In case of *ebpR* the amplification condition was: pre-degeneration at 95 °C for 5 min followed by 35 cycles of degeneration at 95 °C for 1 min, annealing at 52 °C for 30 sec and extension at 72 °C for 50 sec and final extension at 72 °C for 10 min. The amplification products were electrophoresed on 1.5% agarose gel (Photograph-1 and 2). The sequences of primers are provided in table 1.

**Table 1:**
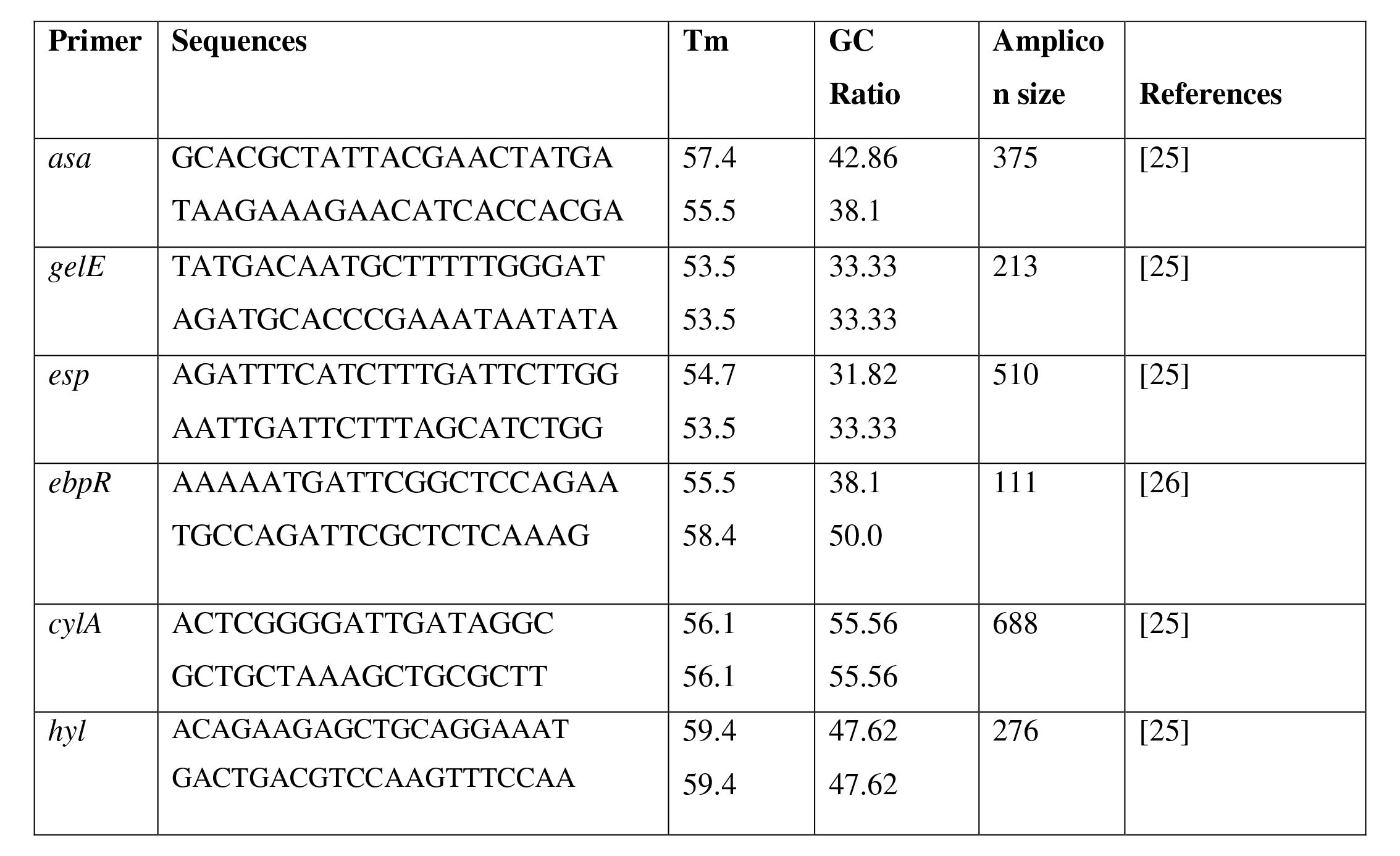
Primer sequences and properties of primers used in the PCR for virulence genes

### Statistical analysis

The association between the biofilm formation, hemolysin production and their encoding genes and their association with antibiotics resistance were evaluated by the Pearson Chi-Square test using SPSS version 22. P values less than 0.05 were regarded as statistically significant.

## RESULTS

### Virulence factors production

Among 87 *Enterococci* isolates, 65(75%) were reconized as *E. faecalis* and 20(23%) as *E. faecium*, 2(2%) as *E. raffinosus*. Table 2 showed among 41 biofilm producer, 34 (52.3%) were *E. faecalis* and 7 (30.8%) were *E. faecium.* Out of 87 isolates, 24 (27.58%) showed hemolysis. Among them 20 (30.8%) were *E. faecalis* and 4 (20%) were *E. faecium.* Gelatin hydrolysis was not detected in any isolates.

**Table 2:**
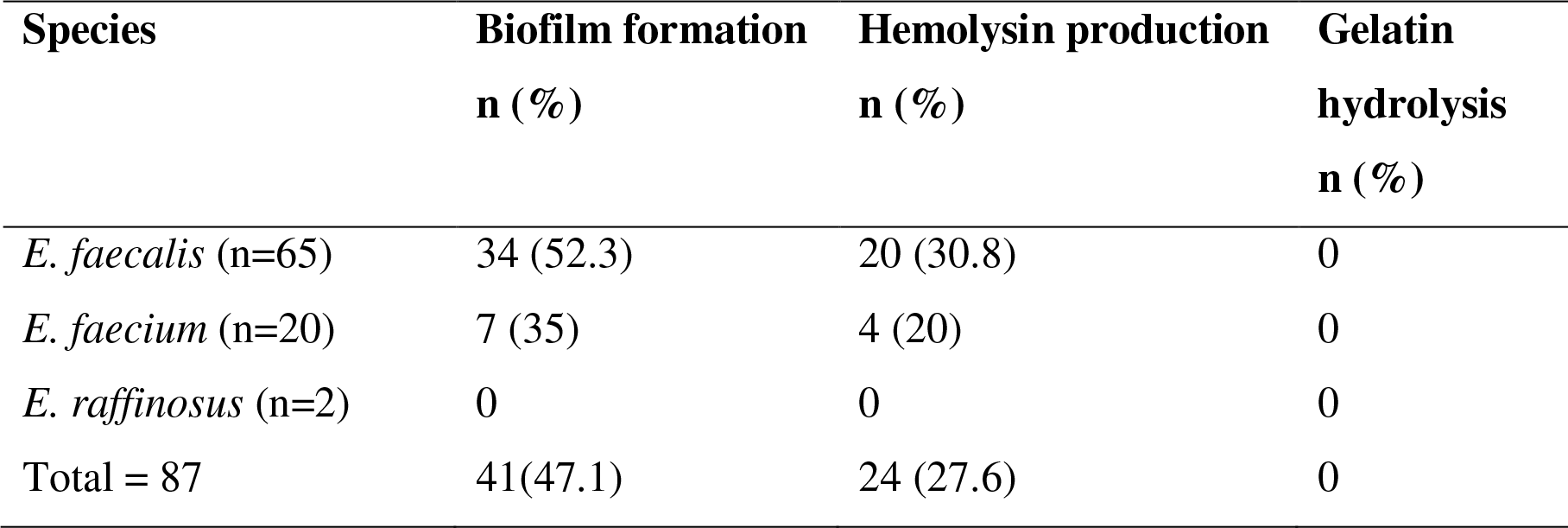
Results of biofilm formation, hemolysin production, gelatin hydrolysis in *Enterococcus* species

| Species | Biofilm formation<br>n (%) | Hemolysin production<br>n (%) | Gelatin<br>hydrolysis<br>n (%) |
| --- | --- | --- | --- |
| *E. faecalis* (n=65) | 34 (52.3) | 20 (30.8) | 0 |
| *E. faecium* (n=20) | 7 (35) | 4 (20) | 0 |
| *E. raffinosus* (n=2) | 0 | 0 | 0 |
| Total = 87 | 41(47.1) | 24 (27.6) | 0 |

### Antibiotic susceptibility testing

Among 87 *Enterococci*, highest resistance were seen in ciprofloxacin 59 (67.8%), gentamicin 53 (60.9%) followed by ampicillin 41 (47.1%). Vancomycin and teicoplanin resistance were detected in 3 (3.44%) isolates. According to the results, linezolid, fosfomycin, quinopristin-dalfopristin were the most effective agents.

### Detection of virulence factors encoding genes

Table 3 and 4 showed the association of biofilm producing encoding genes in biofilm production *in E. faecalis*. The *asa, esp, ebpR* and combination of genes *asa,gel,ebpR*; *asa,esp,gel,ebpR* were found in 65.9%, 64.1%, 60.4%, 87.5%, 86.4% biofilm producing *E. faecalis* which were statistically significant. The *asa, gelE* and combination of genes *esp, hyl, ebpR* were found 100%, *esp* and *ebpR* were found in 85.7 and 50% biofilm producing *E. faecium* which were statistically significant.

**Table 3:** Association of biofilm producing encoding genes and biofilm production in *E. faecalis*

| Virulence gene | Biofilm producer<br>n (%) | Non biofilm producer<br>n (%) | p<br>value |
| --- | --- | --- | --- |
| *asa* only (n=44) | 29 (65.9) | 15 (34.1) | **.001** |
| *esp* only (n=39) | 25 (64.1) | 14 (35.9) | **.02** |
| *gelE* only (n=52) | 28 (53.8) | 24 (46.2) | .619 |
| *ebpR* only (n=53) | 32 (60.4) | 21 (39.6) | **.00** |
| *asa,ebpR* (n=2) | 1 (50) | 1 (50) | .947 |
| *esp,ebpR* (n=1) | 1 (100) | 0 | .396 |
| *esp,ebpR,gelE* (n=2) | 2 (100) | 0 | .1 |
| *asa,esp,ebpR* (n=3) | 3 (100) | 0 | .09 |
| *asa,gelE,ebpR* (n=8) | 7 (87.5) | 1 (12.5) | **.03** |
| *asa,esp,gelE,ebpR* (n=22) | 19 (86.4) | 3 (13.6) | **.000** |

p value was calculated by chai-square test. p value <0.05 was considered statistically significant.

**Table 4:** Association of biofilm producing encoding genes and biofilm production in *E. faecium*

| Virulence gene | Biofilm producer<br>n (%) | Non biofilm producer<br>n (%) | p value |
| --- | --- | --- | --- |
| *asa* (n=2) | 2 (100) | 0 | **.042** |
| *esp* (n=7) | 6 (85.7) | 1 (14.3) | **.000** |
| *gelE* (n=2) | 2 (100) | 0 | **.042** |
| *ebpR* (n=14) | 7 (50) | 7 (50) | **.032** |
| *hyl* (n=4) | 2 (50) | 2 (50) | .482 |
| *esp,ebpR* (n=7) | 5 (71.4) | 2 (28.6) | .012 |
| *asa,gelE,ebpR* (n=1) | 1 (100) | 0 | .162 |
| *asa,esp,gelE,ebpR* (n=1) | 1 (100) | 0 | .162 |
| *esp,hyl,ebpR* (n=2) | 2 (100) | 0 | **.042** |

p value was calculated by chi-square test. p value <0.05 was considered statistically significant.

Table 5 showed the association of *cylA* gene and hemolysin production in *E. faecalis*. Out of 46 *cylA* positive isolates, 20 (43.6%) were detected phenotypically.

**Table 5:**
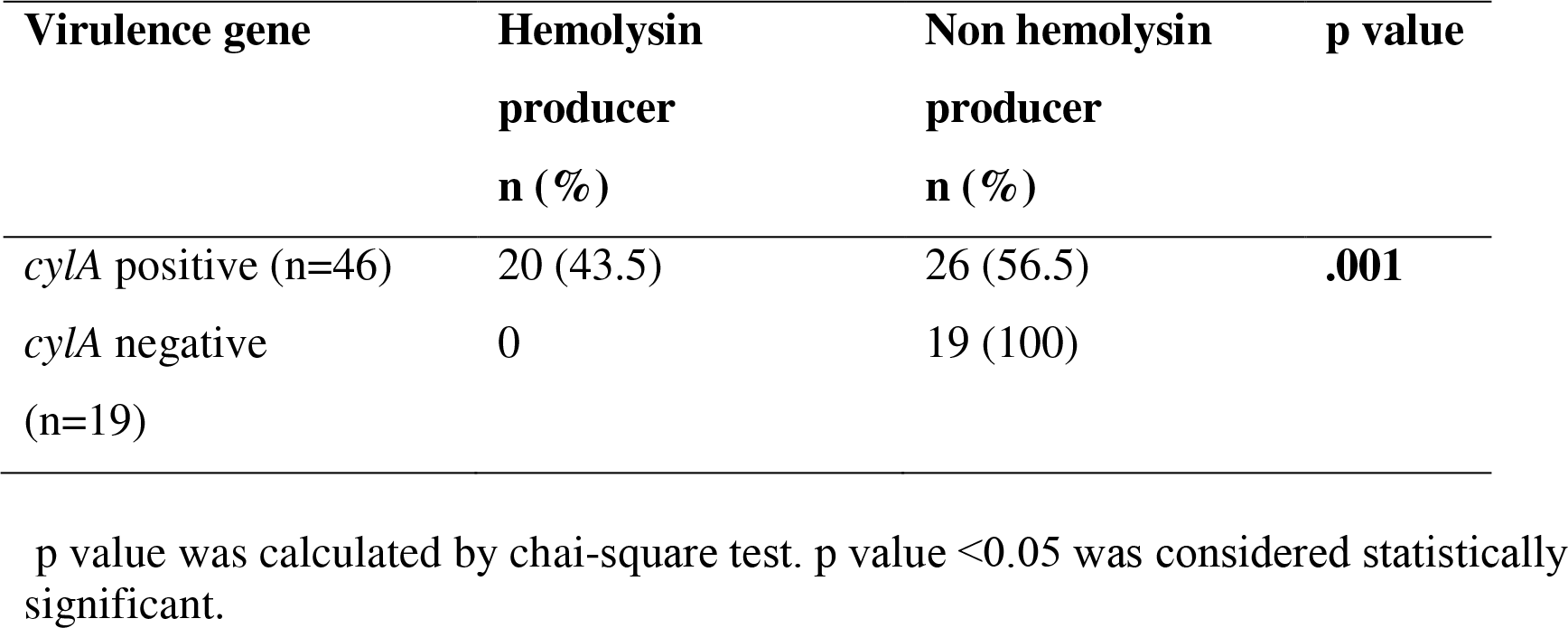
Association of *cylA* gene and hemolysin production in *E. faecalis* isolates

| Virulence gene | Hemolysin<br>producer<br>n (%) | Non hemolysin<br>producer<br>n (%) | p value |
| --- | --- | --- | --- |
| *cylA* positive (n=46) | 20 (43.5) | 26 (56.5) | **.001** |
| *cylA* negative<br>(n=19) | 0 | 19 (100) |  |

p value was calculated by chi-square test. p value <0.05 was considered statistically significant.

**Table 6:** Association of *cylA* gene and hemolysin production in *E. faecium* isolates

| **Virulence gene** | **Hemolysin**<br>**producer**<br>**n (%)** | **Non hemolysin**<br>**producer**<br>**n (%)** | **p value** |
| --- | --- | --- | --- |
| *cylA* positive (n=0) | 0 | 0 |  |
| *cylA* negative (n=20) | 4 (20) | 16 (80) |  |

**Note:** No statistics was computed because *cylA* was a constant.

The association of *cylA* gene and hemolysin production in *E. faecium* isolates were showed in Table None of the hemolysin producing isolate was positive for *cylA* gene. On the other hand, out of 20 *cylA* gene negative isolates, 4 (20%) were hemolysin producers.

### Virulence factors and antibiotic resistance

Biofilm formation and antibiotic resistance of *E. faecalis* and *E. faecium* were shown in table 7 and table 8 respectively. Statistical analysis indicated that there was a significant association between biofilm formation of *E. faecalis* and antibiotic resistance. Significant association was found in case of ampicillin, cotrimoxazole, ciprofloxacin, gentamicin in *E. faecalis*. Antibiotic resistance were higher in biofilm producing *E. faecium* but no significant association of was found.

**Table 7:** Antimicrobial resistance pattern of biofilm and non-biofilm producing *E. faecalis* isolates

| **Antimicrobial agents**<br>**(µg)** | **Biofilm producer**<br>**(34)**<br>**n (%)** | **Non-biofilm producer**<br>**(31)**<br>**n (%)** | **p value** |
| --- | --- | --- | --- |
| Ampicillin (10) | 18 (52.9) | 4 (12.9) | **.001** |
| Cotrimoxazole<br>(1.25/23.75) | 22 (64.7) | 5 (16.1) | **.000** |
| Ciprofloxacin (5) | 29 (85.3) | 11 (35.5) | **.000** |
| Nitrofurantoin (300) | 6 (17.6) | 2 (6.5) | .170 |
| Gentamicin (120) | 24 (70.6) | 9 (29.0) | **.001** |
| Vancomycin (30) | 0 | 0 | - |
| Linezolid (30) | 0 | 0 | - |
| Teicoplanin (30) | 0 | 0 | - |
| Fosfomycin (200) | 0 | 0 | - |
| Quinopristin-dalfopristin (15) | - | - | - |

p value was calculated by chi-square test. p value <0.05 was considered statistically significant.
Note: *E. faecalis* intrinsically resistant to Quinopristin - dalfopristin.

**Table 8:**
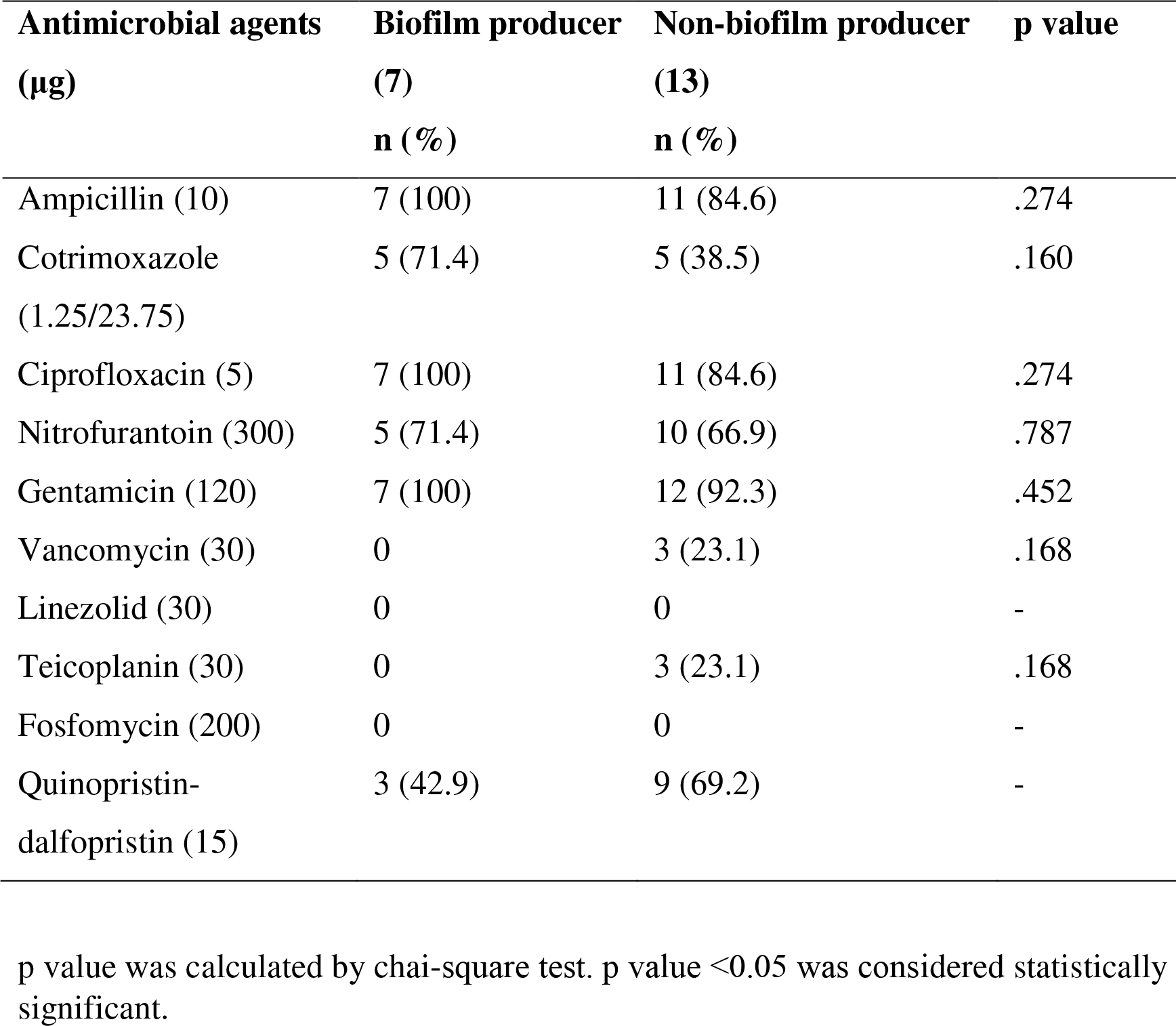
Antimicrobial resistance pattern of biofilm and non-biofilm producing *E. faecium* isolates

Table 9 and 10 showed antibiotic resistance pattern and phenotypic hemolysin formation in *E. faecalis* and *E. faecium.* Significant association was found in case of ampicillin, ciprofloxacin, nitrofurantoin, gentamicin in *E. faecalis*. There was no association of antibiotic resistance found between hemolysin and non hemolysin producing isolates of *E. faecium*.

**Table 9:**
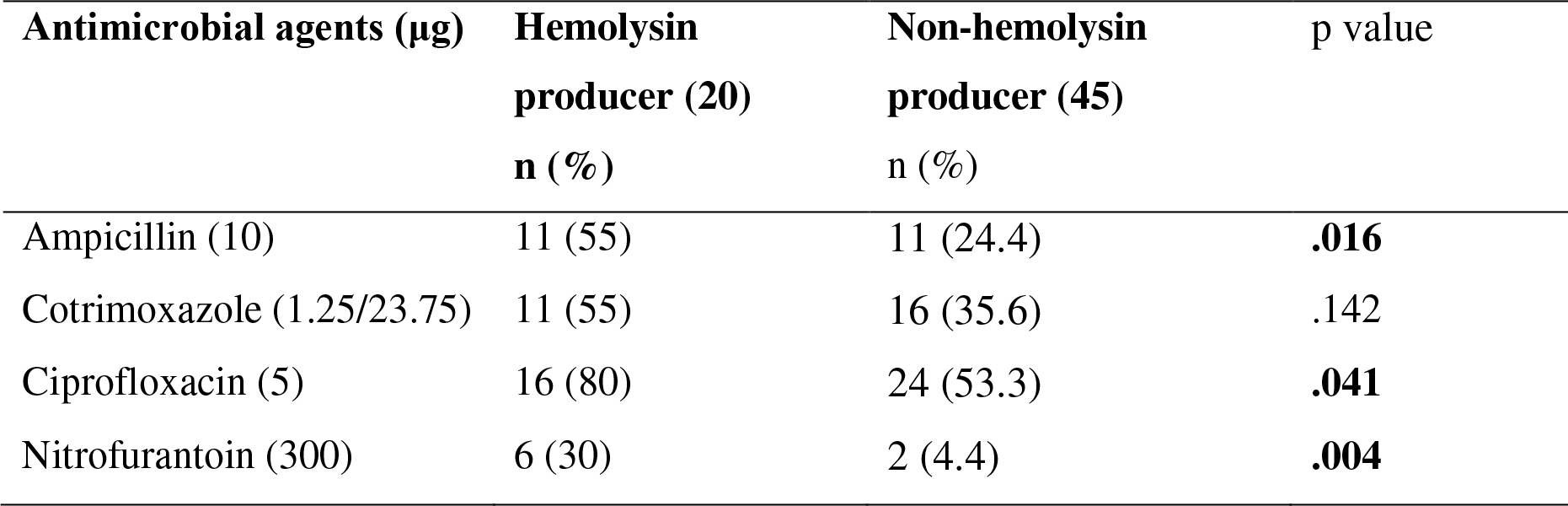
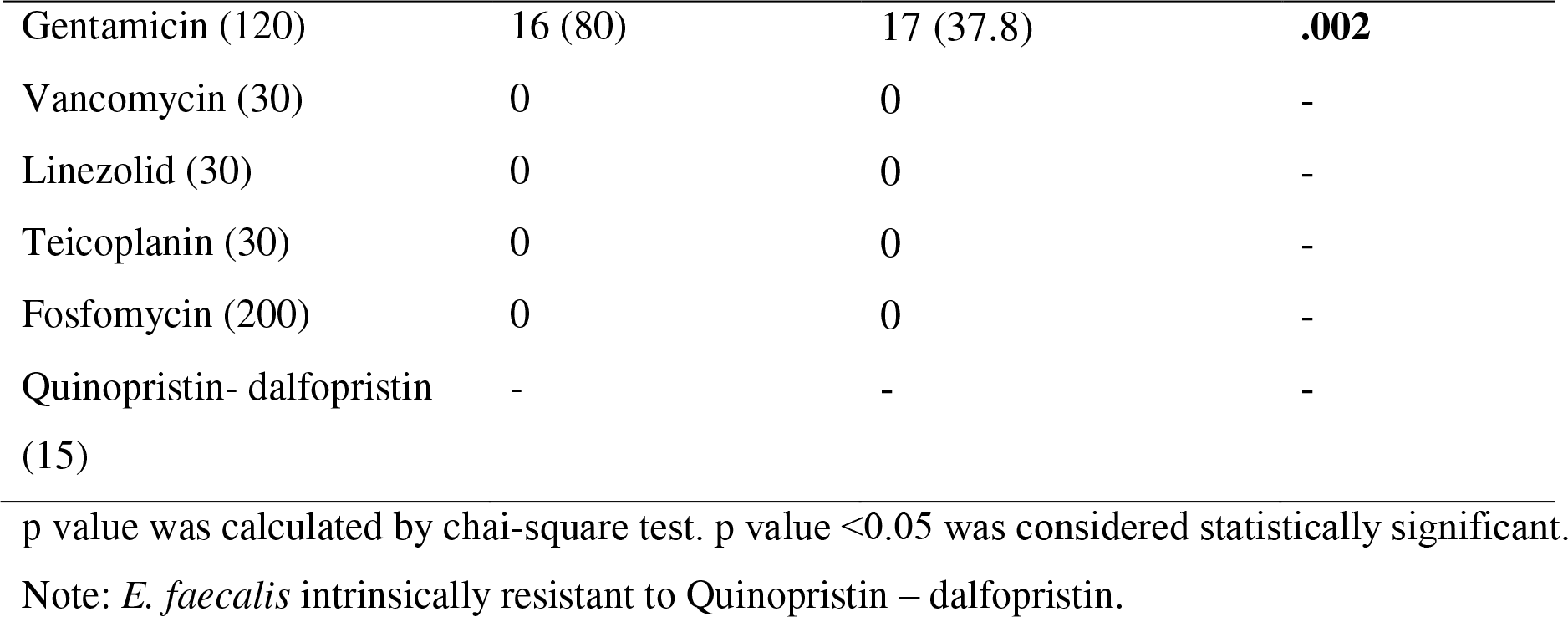
Antimicrobial resistance pattern of hemolytic and non-hemolytic *E. faecalis* isolates

| **Antimicrobial agents (µg)** | **Hemolysin**<br>**producer (20)**<br>**n (%)** | **Non-hemolysin**<br>**producer (45)**<br>**n (%)** | **p value** |
| --- | --- | --- | --- |
| Ampicillin (10) | 11 (55) | 11 (24.4) | **.016** |
| Cotrimoxazole (1.25/23.75) | 11 (55) | 16 (35.6) | .142 |
| Ciprofloxacin (5) | 16 (80) | 24 (53.3) | **.041** |
| Nitrofurantoin (300) | 6 (30) | 2 (4.4) | **.004** |
| Gentamicin (120) | 16 (80) | 17 (37.8) | **.002** |
| Vancomycin (30) | 0 | 0 | - |
| Linezolid (30) | 0 | 0 | - |
| Teicoplanin (30) | 0 | 0 | - |
| Fosfomycin (200) | 0 | 0 | - |
| Quinopristin- dalfopristin (15) | - | - | - |

p value was calculated by chai-square test. p value <0.05 was considered statistically significant.
Note: *E. faecalis* intrinsically resistant to Quinopristin – dalfopristin.

**Table 10:**
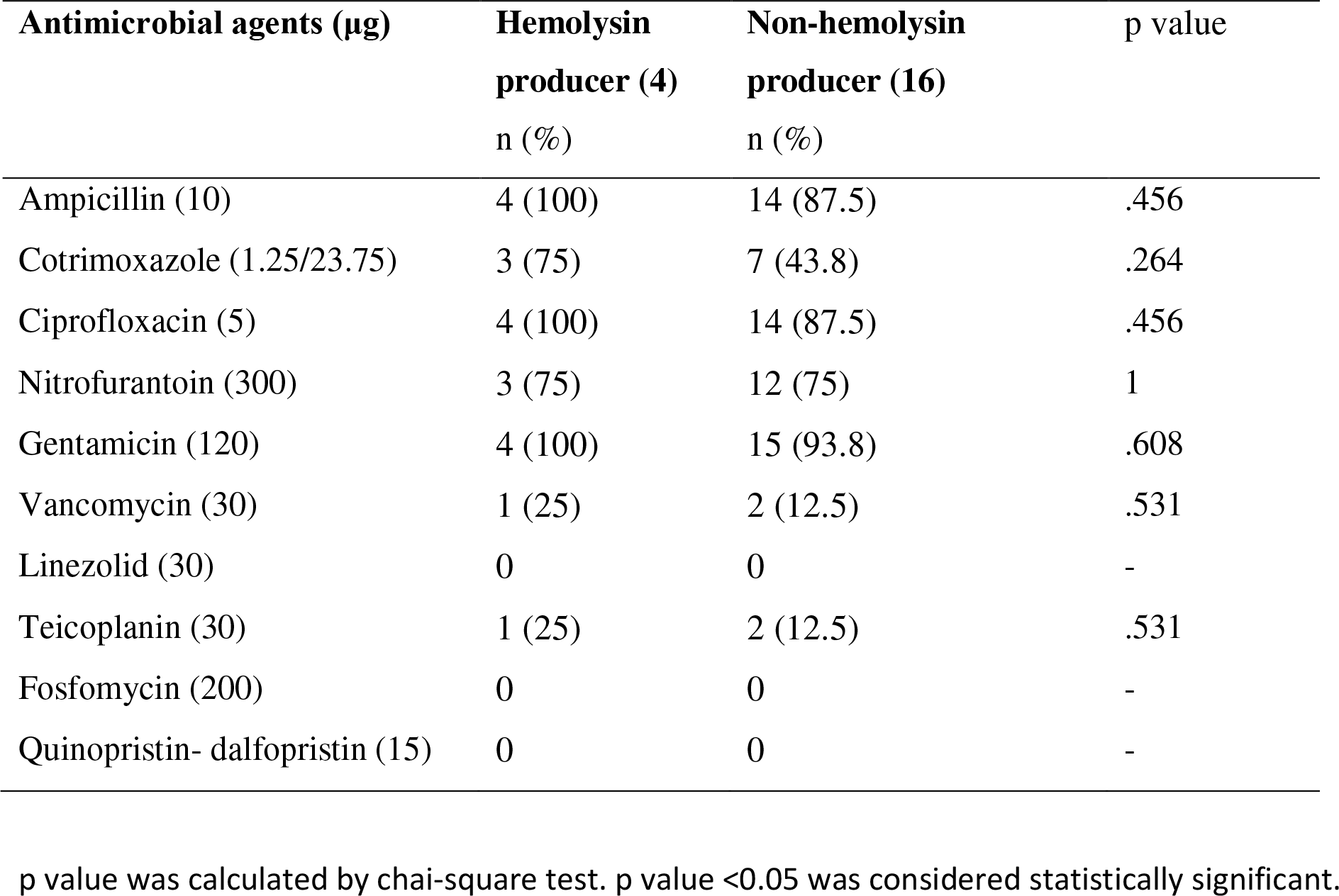
Antimicrobial resistance pattern of hemolytic and non-hemolytic *E. faecium* isolates

| Antimicrobial agents (µg) | Hemolysin<br>producer (4)<br>n (%) | Non-hemolysin<br>producer (16)<br>n (%) | p value |
| --- | --- | --- | --- |
| Ampicillin (10) | 4 (100) | 14 (87.5) | .456 |
| Cotrimoxazole (1.25/23.75) | 3 (75) | 7 (43.8) | .264 |
| Ciprofloxacin (5) | 4 (100) | 14 (87.5) | .456 |
| Nitrofurantoin (300) | 3 (75) | 12 (75) | 1 |
| Gentamicin (120) | 4 (100) | 15 (93.8) | .608 |
| Vancomycin (30) | 1 (25) | 2 (12.5) | .531 |
| Linezolid (30) | 0 | 0 | - |
| Teicoplanin (30) | 1 (25) | 2 (12.5) | .531 |
| Fosfomycin (200) | 0 | 0 | - |
| Quinopristin- dalfopristin (15) | 0 | 0 | - |

p value was calculated by chai-square test. p value <0.05 was considered statistically significant.

## DISCUSSION

Out of 87 *Enterococci* isolates, *E. faecalis* 65 (75%) and *E. faecium* 20 (23%) were most common. *E. raffinosus* was also isolated 2 (2%) in this study. Biofilm-producing *Enterococci* are responsible for recurrent, chronic and antibiotic-resistant infection [9]. In present study, about 41 (47.1%) *Enterococci* were biofilm producers. Sixty eight percent and 21.9% biofilm producing *Enterococci* were also reported by Sindhanai *et al* and Shridhar *et al* respectively [27, 28].

In this study *E. faecalis* produced more biofilm than *E. faecium*. Thirty four (52.3%) *E. faecalis* and 7 (35%) *E. faecium* were biofilm producer in this study. This results were similar to the study conducted by Zheng *et al* found 47.2% *E. faecalis* produced biofilm [29]. This findings were also supported by other studies done by Banerjee & Anupurba *et al* where they found 27.09% *E. faecalis* and 25.16% *E. faecium* isolates produced biofilm respectively [30].

Weng *et al* reported that *E. faecium* produce more biofilms than *E. faecalis* (59.3% versus 49.0%) [31]. The conflict may be due to number of *E. faecium* in that study and geographic differences.

In present study, about 24 (27.6%) hemolysin producing *Enterococci* was observed. Similarly, 31.61%, 18.25%, 15% hemolysin producing *Enterococci* were reported by other study [30, 20, 32]. Kashef *et al* reported 58% hemolysin producing *Enterococci* which was quite higher than this study [33]. None of the *Enterococcus* species was positive for gelatin hydrolysis which was similar to the finding of a study done in Egypt by Hashem *et al* [34]. Manavalan *et al* reported that 19.84% *Enterococci* isolates were gelatinase producers which was dissimilar to the finding of this study [20]. Kashef *et al* also showed that many *gelE* positive isolates failed to secrete gelatinase in their study [33]. Gelatinase encoded by *gelE* gene and is regulated by the transmembrane protein, FsrB. It is controlled by locus *fsr,* deletions within locus *fsr* produce mutants that do not synthesize gelatinase [35, 36].

In this study, hundred percent of biofilm producing *E. faecium* and 65.9% biofilm producing *E. faecalis* isolates had *asa* gene which was statistically significant (p value .042 and .001 respectively). Banerjee & Anupurba *et al* found the *asa* gene was associated with biofilm formation which is similar to our findings but the results was dissimilar to the finding by Fallah *et al* [30, 26].

The *esp* gene was found in 64.1% and 85.7% of biofilm producing *E. faecalis* and *E. faecium* in this study respectively which were statistically significant (p value .02 and .000 respectively). Toledo-arena *et al* found a correlation between the *esp* gene and biofilm formation [22]. Other studies suggest that the *esp* gene is not necessary nor sufficient for the biofilm production in *E. faecalis* and *E. faecium* [37, 38]. In this study, 53.8% and 100% biofilm producing *E. faecalis* and *E. faecium* had *gelE* gene which was not statistically significant (p value 0.619) for *E. faecalis* but for *E. faecium* (p value 0.042). Other study found that *gelE* gene was associated with biofilm formation in *Enterococci* which supported our findings in case of *E. faecium* [30]. Other studies suggest that gelatinase was not required for biofilm formation [39].

In this study, 60.4% and 50% biofilm producer of *E. faecalis* and *E. faecium* had *ebpR* gene which was statistically significant (.006, .032 respectively). Kafil and Mobarez *et al* reported that 93.6% biofilm producers had *ebp* gene which is almost similar to this study [7].

All biofilm producing *E. faecalis* and *E. faecium* isolates of this study had multiple biofilm forming genes. In present study, combination of *asa, gelE, ebpR* genes and *asa, esp, gelE, ebpR* genes had significant association with biofilm formation (p value .03 and .000 respectively) in *E. faecalis*. Combination of *esp, ebpR* and *esp, hyl, ebpR* genes had significant association with biofilm formation (p value .012 and .042 respectively) in *E. faecium*. This was supported by other studies where they found that individual biofilm related genes such as *esp, asa, ebpR, gelE* did not appear to be sufficient for biofilm formation in *Enterococci* [26, 40]. On the contrary, other study reported that single biofilm forming gene was associated with biofilm formation [30]. Biofilm formation is complex process and depends on various factors in *Enterococcus* strains [41].

Cytolysin plays an important role in the severity of human infections [42, 43]. In present study, hemolytic activity was detected phenotypically in 30.8% *E. faecalis* where *cylA* was present in 43.5% which was statistically significant (.001). Safari *et al* showed that 41% *Enterococci* carried the *cylA* gene and hemolytic activity was detected in 38% of *cylA* positive isolates which supports the findings of this study [40]. This lack phenotypic/genotypic expression of cytolysin might suggest the missing genes in the *cyl* operon [44].

The *cylA* gene was absent in all (4) hemolysin producing isolates of *E. faecium* in this study. DeVuyst *et al* reported that the cytolysin structural gene was always present in β-hemolytic *E. faecalis* while this gene could not be detected in *E. faecium* [45]. So, the hemolysis of *E. faecium* must be caused by another cytotoxic component. In other study, 75 isolates with hemolytic activity were negative for *cylA* gene suggesting possible role of other genes in hemolytic activity which was similar to the finding of this study in *E. faecium* isolates [46].

Treatment of biofilm forming *Enterococci* are difficult because they are more antibiotic resistant. [30]. Regarding antibiotic resistance, higher resistance was observed in biofilm producers compared to non biofilm producers of *E. faecalis*. Resistance to some antibiotic including ampicillin (52.9% vs. 12.9%), cotrimoxazole (64.7% vs. 41.5%), ciprofloxacin (85.3% vs. 35.5%), HLGR (70.6% vs. 29.6%) were significantly higher in biofilm producer than non biofilm producers. Similar results were reported by Fallah *et al* and they found significantly higher resistance among biofilm positive isolates [26]. Resistance to antibiotics were higher in biofilm producing strains of *E. faecium*. Biofilm producers were 100% resistance to ampicillin, ciprofloxacin, HLGR followed by 71.4% resistance to cotrimoxazole and nitrofurantoin which were higher than non biofilm producing isolates. Sieńko *et al* also reported that biofilm producing *E. faecium* strains were more antibiotic resistant [47].

Hemolysin producing *E. faecalis* isolates of this study were more antibiotic resistant than non hemolysin producing isolates. Resistance to some antibiotic including ampicillin (55% vs. 24.4%), cotrimoxazole (55% vs. 35.6%), ciprofloxacin (80% vs. 53.3%), HLGR (80% vs. 62.2%), nitrofurantoin (30% vs. 4.4%) were higher in hemolysin producer than non hemolysin producers in this study. Sachan *et al* reported significant relation was found between hemolysin production and HLGR which supports this study [48].

In this study, hemolysin producing isolates of *E. faecium* were more antibiotic resistant than non hemolysin producing isolates. Resistance to some antibiotic including ampicillin (100% vs. 87.7%), cotrimoxazole (75% vs. 43.8%), ciprofloxacin (100% vs. 87.5%), HLGR (100% vs. 93.8%) were higher in hemolysin producer than non hemolysin producers but no significant association was observed which supports the finding of a study by Jankoska *et al* [49]. High number of resistance was also seen in non hemolysin and non biofilm producing isolates of *E. faecium*. The reason may be due to *E. faecium* is more drug resistant.

## CONCLUSION

This study reveals the ability of *Enterococci* to produce biofilm formation and hemolysin production. The biofilm encoding genes *ebpR, asa, esp, gelE* found in biofilm producing isolates of both species and hemolysin encoding gene *cylA* in hemolysin producing *E. faecalis* isolates suggests potential link between these genes in biofilm formation and hemolysin production. Linezolid and fosfomycin, quinoristin - dalfopristin remains the most effective antimicrobial agent. This trend of multidrug resistance in *Enterococci* is a matter of concern.

## ACKNOWLEDGEMENT

The corresponding author is thankful to the Departmentof Microbiology and Immunology, Bangabandhu Sheikh Mujib Medical University (BSMMU) for providing necessary facilities to carry out the research.

## ETHICAL APPROVAL

The study was ethically approved by Institutional Review Board (IRB) of Bangabandhu Sheikh Mujib Medical University, Dhaka, Bangladesh [NO.BSMMU/2019/8188, Date-29/07/2019]

## AUTHOR CONTRIBUTION

Abu Naser Ibne Sattar was responsible for the conception of the study. Abu Naser Ibne Sattar, Rehana Razzak Khan, Fatima Afroz, Nahidul Islam, Sharmin Chowdhury, Tahani Momotaz and Mohammad Tanvir Sarwar participated in its design and coordination. Tahani Momotaz was chief investigator and responsible for the acquisition, analysis and interpretation of the data. Mohammad Tanvir Sarwar, Abu Naser Ibne Sattar and Rehana Razzak Khan reviewed the results and statistical analyses. Tahani Momotaz drafted the manuscript and all the authors contributed substantially to its revision. All authors met ICMJE authorship criteria and have read and approved the final manuscript.

---

**Figure 1:**
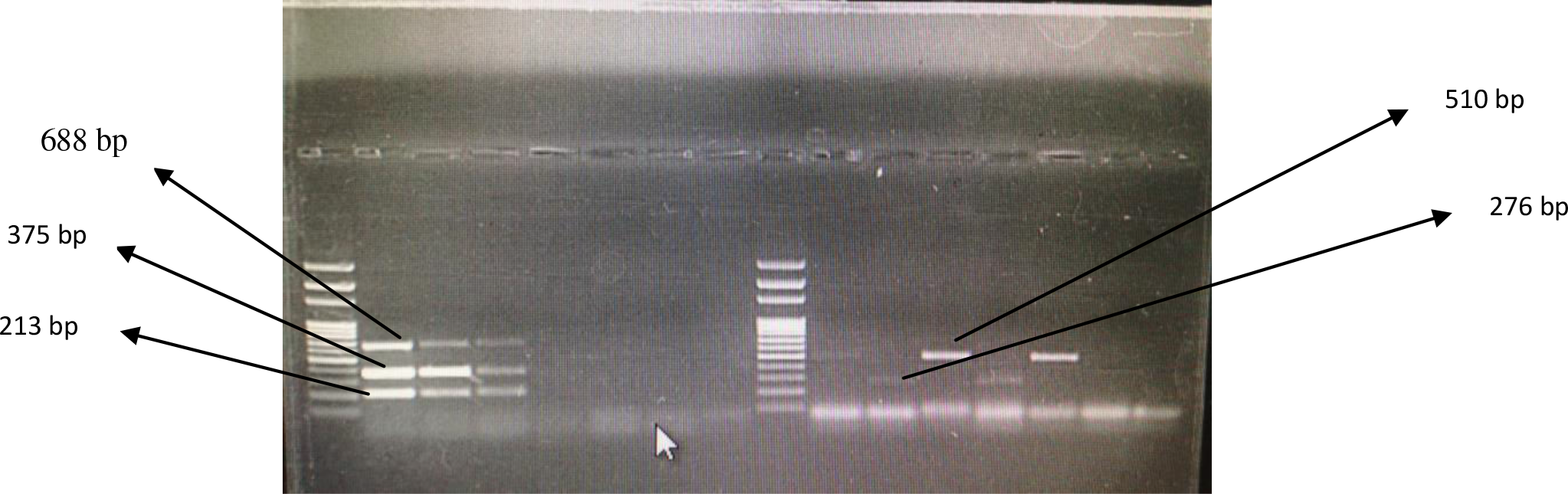
Agarose gel electrophoresis analysis showed amplified DNA product of *asa* (375 bp), *gelE* (213 bp), *cylA* (688), *esp* (510 bp), *hyl* (276 bp) by multiplex PCR. Lane M: DNA ladder (100 bp), Lane1: positive control of *asa* (375 bp), *gelE* (213 bp)*, cylA* (688 bp) genes, Lane 2, 3: *asa, gelE, cylA* positive *Enterococci*, Lane 4, 5, 6: *asa, gelE, cylA* negative *Enterococci*, Lane 7: negative control of *asa, gelE, cylA* gene, Lane 9: positive control of *hyl* gene (276 bp), Lane 10: positive control of *esp* gene (510 bp), Lane 11: *hyl* positive *Enterococci,* Lane 12: *esp* positive *Enterococci,* Lane 8, 13: *esp, hyl* negative *Enterococci*, Lane 14: negative control of *hyl, esp* gene.

**Figure 2:**
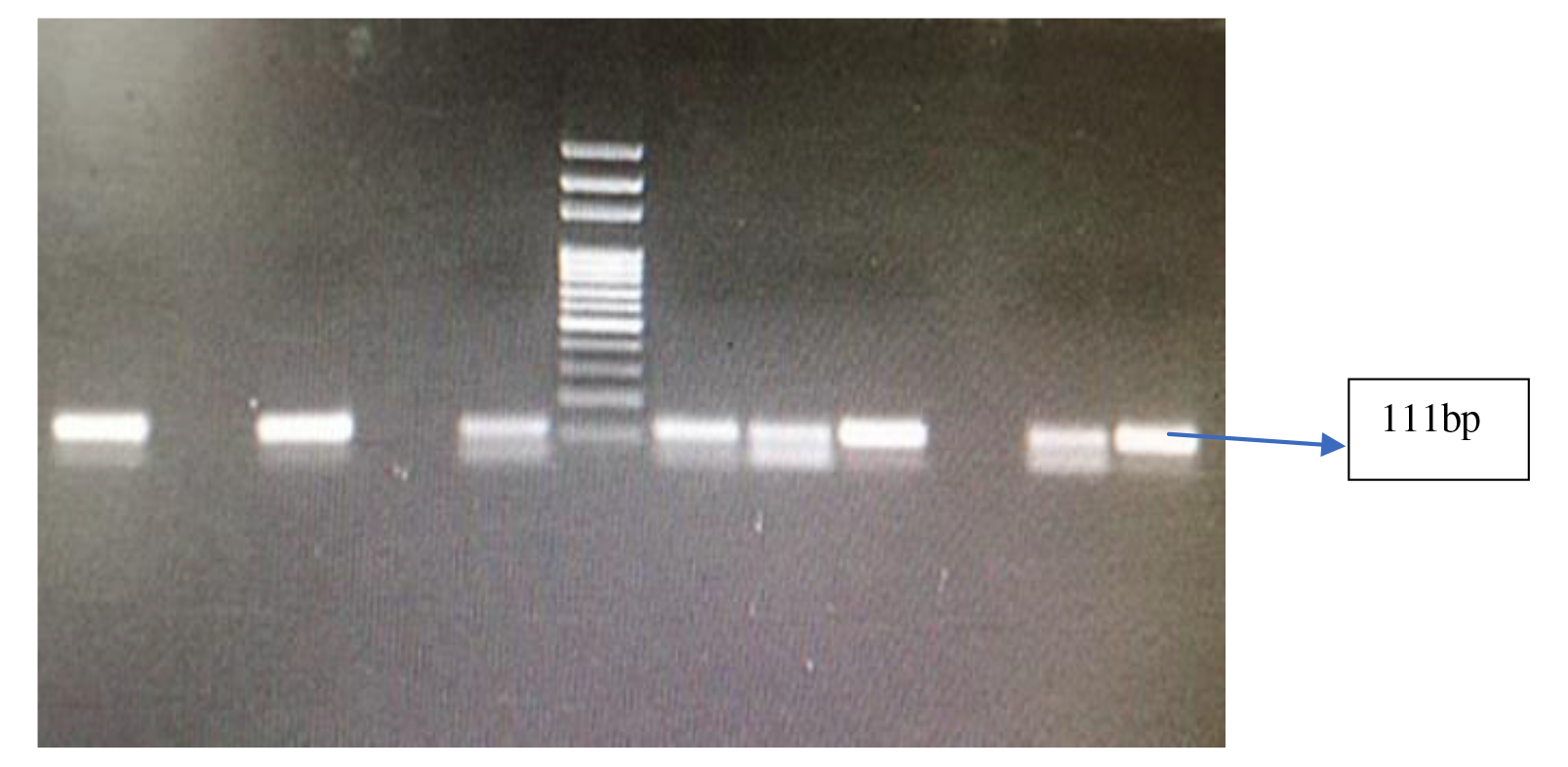
Agarose gel electrophoresis analysis showed amplified DNA product of *ebpR* (111bp). Lane M: DNA ladder (100 bp), Lane 1: positive control of *ebpR* (111 bp) gene, Lane 2: negative control of *ebpR* gene, Lane 3,5,6,7,8,10 *ebpR* positive *Enterococci*, Lane 4, 9: *ebpR* negative *Enterococci*.

